# Water-immersion finger-wrinkling improves grip efficiency in handling wet objects

**DOI:** 10.1101/2020.11.07.372631

**Authors:** Nick J. Davis

## Abstract

For most people, immersing their hands in water leads to wrinkling of the skin of the fingertips. This phenomenon is very striking, yet we know little about why it occurs. It has been proposed that the wrinkles act to distribute water away from the contact surfaces of the fingertip, meaning that wet objects can be grasped more readily. This study examined the coordination between the grip force used to hold an object and the load force exerted on it, when participants used dry or wrinkly fingers, or fingers that were wet but not wrinkly. The results showed that wrinkly fingers reduce the grip force needed to grip a wet object, bringing that force in line with what is needed for handling a dry object. The results suggest that enhancing grip force efficiency in watery environments is a possible adaptive reason for the development of wrinkly fingers.

## Background

When human hands and feet are immersed in water, over time the skin becomes wrinkled. The wrinkling is mainly confined to the pads of the fingertips and to the toes. Explanations for the wrinkling of the skin include a passive response of the skin to immersion, or an active process that creates the wrinkles for a functional purpose. There is overwhelming evidence that finger-wrinkling is an active process. The small blood vessels of the fingertip constrict, which creates valleys in the skin surface, triggered by water entering sweat pores (1). Note that a passive explanation would usually assume that water absorbs into the skin, pushing up ridges. This vasoconstriction appears to occur most readily at a temperature of around 40° Celsius, or the temperature of a warm bath (2). People with autonomic neurological conditions including Parkinson’s, cystic fibrosis, congestive heart failure or diabetic neuropathy may show abnormal or asymmetric wrinkling in the affected parts of the body (3–5).

Given that finger-wrinkling is actively maintained, the natural question is why this would happen. It has been suggested that active finger-wrinkling is an adaptation to aid grasping of objects in watery environments. In order to grasp an object, the grip force used to stabilise the object must be enough to balance the load force, which is generated by the mass of the object and is affected by movements of the object, and must take into account the friction of the interface between the fingertips and the object surface (6–8). Put simply, a wet stone needs to be gripped harder than the same stone when it is dry, as the friction of the contact surface is reduced due to the water. Many authors have linked the wrinkling of the fingertips to this grip- and load-force coordination, with the suggestion that the wrinkles act in the same way as the treads on a car tyre, which help to channel water and to provide ridges of drier contact surfaces on the road (9).

If finger wrinkles do indeed aid grasping, we would expect to see this reflected in the grip force used to manipulate an object. Grip and load force are tightly coupled in both static and dynamic grasps. In consciously-initiated movements grip force changes in parallel with the change in load, and slightly precedes it, suggesting a degree of planning of grip force to cope with the changes of load induced by the inertia of the grasped object (6).

In this study grip and load force was measured in a task where participants gripped an instrument between finger and thumb, and used this to track a load force target as it moved across a screen. It was hypothesised that participants with wrinkly fingers would be more efficient in their grip force than participants with wet but non-wrinkly fingers.

## Methods

### Ethics

All data reported here were collected while the author was in residence at the Science Museum in London, UK. Ethical approval was granted both by the author’s then institution, the Department of Psychology, Swansea University, UK, and by the Science Museum. Participants aged 18 or over gave written informed consent to take part in the study, while parental consent was given in the case of people under 18.

### Participants

After giving informed consent, participants allocated themselves to one of three conditions: these were ‘Dry’ for people who used dry fingers when taking part, ‘Wet’ for people who briefly dipped their fingers in water prior to data collection, or ‘Wrinkly’ for people whose fingers were wrinkled during the experiment. 546 people initially took part in the experiment.

### Procedure

To generate wrinkled fingers, participants immersed their preferred hand in a bath of water kept at 30° Celsius, until the fingertips were visibly wrinkled to the satisfaction of the experimenter. To collect grip- and load-force data, two load cells were linked together such that the participant gripped one load cell between the finger and thumb of their preferred hand (NovaTech F255, NovaTech Measurements Ltd., UK), and could push or pull the second load cell vertically (NovaTech F256). The arrangement of the load cells is shown in Figure 1. The load cells were connected to a laptop that displayed the output of the vertical load, using a custom program written in Matlab (The Mathworks, Natick, MA). During a trial, participants were asked to follow a trace that appeared on the screen of the laptop. The target trace appeared as a solid blue line that swept left-to-right across the screen, and the instantaneous output of the vertical load cell was shown as a red circle, with the ‘history’ of this vertical force shown on the screen as pale dots. Each trial lasted 15 sec. The target trace was static at 0.5 N for 3.5 sec, then rose to 2 N over the course of 3 sec, then was static at 2 N for 4 sec, and dropped to 0.5 N over 3 sec, where it remained for the rest of the trial. Participants each contributed eight trials. Data from the both load cells were digitised at 1000 Hz and stored for later analysis.

**FIGURE 1.**
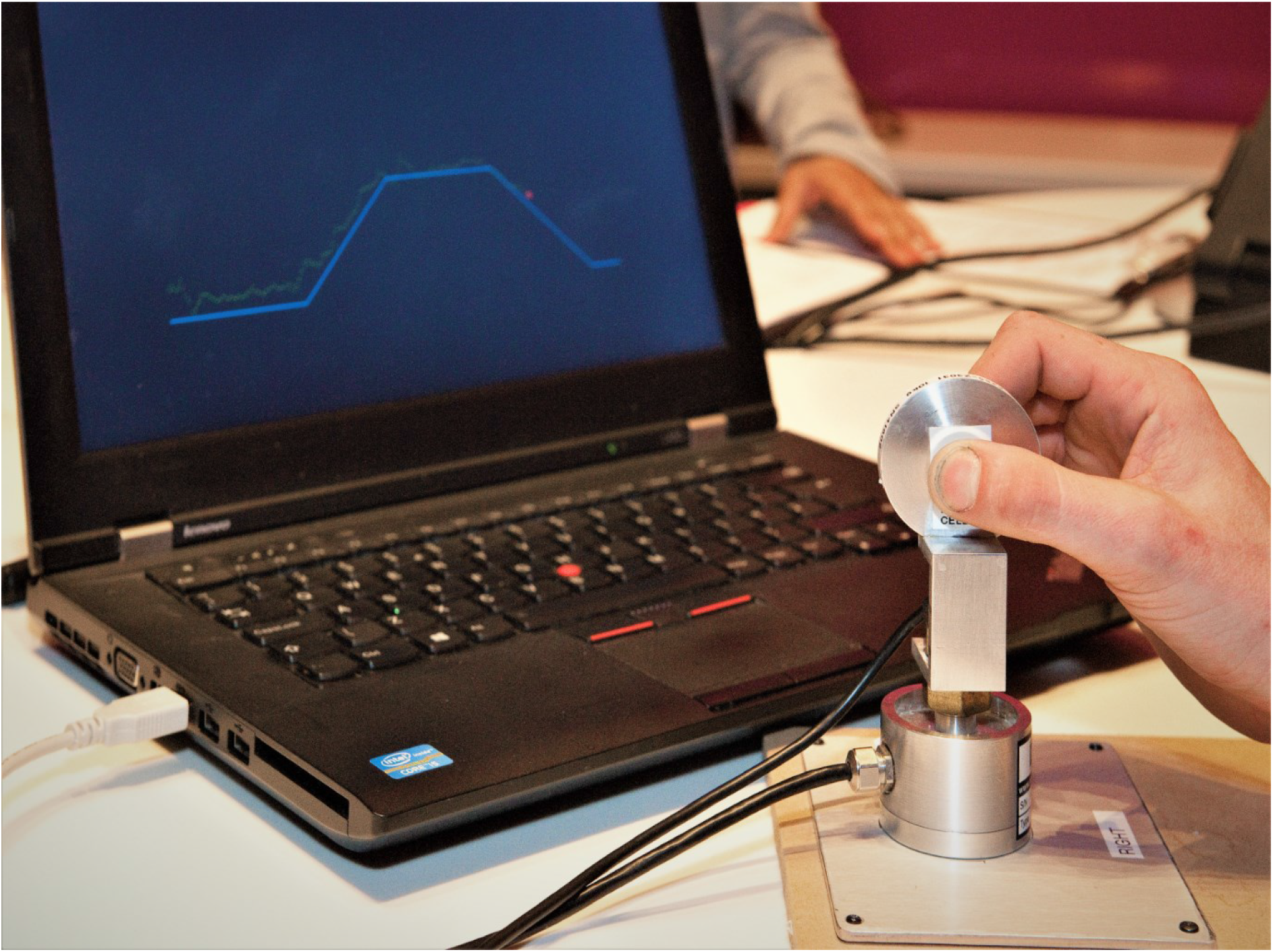
Picture of the equipment in use. The participant is gripping a load cell between finger and thumb. The participant’s task is to pull up on the second load cell to match a force trace that appears on the laptop monitor. The current load force is shown as a red circle, and the history of the participant’s force is shown as a trail of green dots.

### Data analysis

The grip- and load-force data and the target trace were aligned in time, and the load cell data were low-pass filtered with a second-order Butterworth filter set at 20 Hz. Task performance was assessed by determining the correlation between the target force and the load trace, and subjecting these values to a one-way analysis of variance between groups. The primary measure of interest was the ratio of the grip force to the load force. A segment of 3,000 samples (3 sec) was taken from the static phase of the lift. The mean load force and the mean grip force were taken from this time range, and a mean grip:load force ratio was taken for each participant. These measures were subjected to a one-way analysis of variance, with fingertip condition as a between-subjects factor (Wet, Dry or Wrinkled). The lag between the change of grip force and the change of load force was also measured, using a cross-correlation between the two traces with a maximum lag of ±150 ms. Individual trials were excluded from analysis if the load force trace did not significantly differ from 0 N in the second half of the static hold (suggesting that the participant was not following the target), or if the grip force was more than ten times greater than the load force (suggesting an excessively high grip), and a participant was excluded from analysis if more than three trials were excluded based on these criteria.

### Data accessibility

All raw and processed data files from this experiment are available at Figshare.com: https://doi.org/10.6084/m9.figshare.13201169. All analyses were conducted in Matlab, and the analysis routine is available in the same repository.

## Results

After automatic analysis of the force traces, 516 participants’ data were analysed. Of these participants, 309 identified as female and 217 as male, and the mean age was 17.7 (SD 13.1). 55 participants chose to use their left hand and 461 their right. 231 participants chose to take part in the Dry condition, 74 in the Wet, and 211 in the Wrinkly condition.

Figure 2 shows the mean traces for the three different conditions. The participants’ target force is shown as a black line. The load force traces follow this target line reasonably well, which was expected as the load force was visible to participants as a cursor. The grip force exceeds the load force, as expected. However there is a clear separation between the three traces, with participants with wet fingers using more fingertip force than those who used dry fingers, and with the wrinkly fingers lying between the two.

**FIGURE 2.**
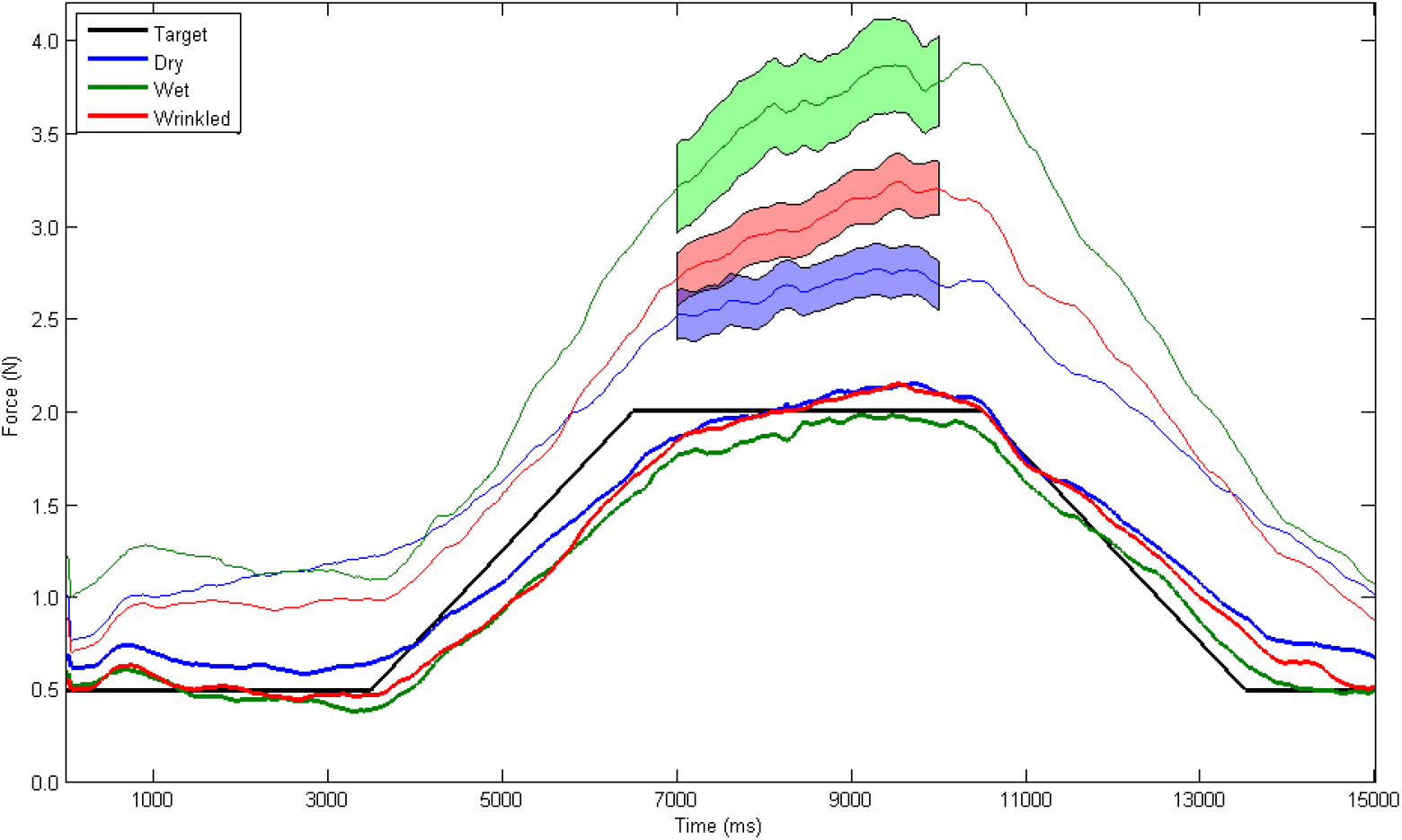
Mean grip force (thinner traces) and load force (thicker traces) when participants tracked a load weight target (black line). Participants with wrinkled fingers produced a grip force that did not differ from that used by people with dry fingers in the static hold phase (indicated with ±1 standard error), however people with wet but non-wrinkly fingers used a significantly higher amount of grip.

The correlation between the participants’ load force and the target force was good, with no differences between groups [F(2,513)=0.953, p=0.386], and with a mean Pearson’s correlation coefficient of 0.628, suggesting that the participants’ primary task was executed successfully. A one-way analysis of variance found that the mean of the ratio of grip to load force was different between the conditions [F(2,513)=8.136, p=0.0003]. Post hoc comparisons found that the ratio was significantly higher for Wet than for Dry (p=0.0002) or for Wrinkly (p=0.0063), but that Dry and Wrinkly did not differ from each other (p=0.384). There was a small but significant correlation of this ratio with the age of the participant [r(514)=−0.149, p<0.001]. The ratio declined by 0.014 per year of age, although the variance explained by a linear regression was very low (R^2^=0.022).

The lag between the change in grip force and the change in load force was not significantly different between the three groups of participants [F(2,513)=0.359, p=0.699], but the overall lag was significantly different from simultaneity, with GF leading LF by 22.62 ms. The lag between grip and load force declined significantly with age [r(514)=−0.338, p<0.001], with the lead of grip over load change declining by 1.36 ms for each year of age, although the variance explained by a linear regression of these values was low (R^2^ = 0.114).

## Discussion

There is now converging evidence that finger-wrinkling is an adaptation that aids object manipulation in wet environments (9). This study has shown that grip efficiency, or the amount by which grip force exceeds the load exerted by the object, is improved when a person has wet and wrinkly fingers, compared to when their fingers are wet but not wrinkly. This ratio of grip force to load force is not significantly different between wrinkly and dry fingers, nor does the relative time difference between the rise of grip force and the rise of load force. Both the grip-to-load ratio and the time difference correlated weakly but significantly with the participants’ age.

Grip and load force coordination is an important aspect of object handling. The ability to generate the correct amount of grip force for a given load means that the minimum necessary amount of energy is used by the muscles that control the fingers and hands, and means that objects are less likely to be dropped or to be crushed. Efficient grip force coordination is seen in many extant primates, and is likely to have evolved early in the primate lineage (10). The grip force required to stabilise a wet object is greater than the force required for a dry object, since the coefficient of friction of the digit-object interface is reduced (8). It would therefore benefit an animal to gain an advantage in handling wet objects, as this would increase success in hunting and foraging in watery environments. Fingertip wrinkles would seem to afford such an advantage in object handling, and may plausibly aid travel and clambering in wet areas, especially if combined with wrinkled toes.

A previous study of object manipulation with wrinkled fingers found that wet objects were moved more quickly when the fingers were wrinkly compared to dry (11). Importantly, there is no difference in tactile sensitivity in wrinkled fingers compared to dry (12), meaning that people are not trading off acuity for friction at the fingertip. It is therefore reasonable to wonder why healthy people do not have permanently wrinkled fingers. The answer presumably lies in the changes in the mechanical properties of the finger tissues, where there may be lower shear resistance when the finger is wrinkled (13). Previous studies have also suggested differences in manipulation across the lifespan (14–16); the present results show age-related effects, although they are rather weak in this sample.

The results presented here should be read in the context of the experiment itself. The age distribution in this sample was rather low, reflecting the public engagement setting of the data collection. Although the effects of age in the data reported here were very small, they were nevertheless statistically significant, so this should be taken into account when comparing these results with others. There may also be effects on performance from inter-individual differences in hand size, in levels of subcutaneous fat, or in lifestyle or genetic factors that were not measured here. Finally the experiment only tested one target force pattern and one fingertip contact surface; it is likely that changing the dynamics of the load and the properties of the object would affect grip force coordination (6, 17).

In summary, this experiment investigated fine motor coordination when the fingers are affected by water-induced finger-wrinkling. Finger-wrinkling improves grip force coordination when compared to fingers that are wet but not wrinkly, and brings the performance to a level comparable with dry fingers. These results help to explain why humans and their close primate relatives may have developed finger-wrinkling as an adaptation to aid locomotion and foraging in wet environments.

## Acknowledgements

The author is grateful to the staff of the Science Museum, London, for access to the Live Science gallery. Particular thanks are due to Georgie Ariaratnam, and to the many visitors who took part in, or discussed, the experiment.

